# Novel monoclonal antibodies targeting distinct sites on placental binding *P. falciparum* antigen VAR2CSA synergistically enhance parasite phagocytosis

**DOI:** 10.64898/2026.04.28.721320

**Authors:** Vivin R. Kokuhennadige, Christopher A. Gonelli, Oscar H. Lloyd Williams, Robyn Esterbauer, Andrew Kelly, Wina Hasang, Holger W. Unger, Paula Tesine, Alice Mengi, Benishar Kombut, Ching-Seng Ang, Adam K. Wheatley, Stephen J. Rogerson, Elizabeth H. Aitken

## Abstract

Placental malaria, due to the sequestration of *Plasmodium falciparum*-infected erythrocytes (IE), causes adverse pregnancy outcomes. The sequestration is mediated by VAR2CSA, a protein that binds to placental chondroitin sulfate A (CSA). VAR2CSA antibodies protect against adverse pregnancy outcomes; however, no licensed VAR2CSA-based vaccine or therapeutic exists to date. We identified and expressed VAR2CSA-specific IgG1 monoclonal antibodies (mAbs) using B cells of malaria-exposed Papua New Guinean women. VAR2CSA mAbs were characterised by their ability to recognise eight heterologous CSA-binding *P. falciparum* strains, to neutralise CSA binding and/or to induce phagocytosis of IEs by THP-1 monocytes. We identified 16 mAbs, all of which targeted just two of the six domains of VAR2CSA, DBL3X and DBL5ε. Cross-reactivity varied between mAbs, but was highest among mAbs to DBL5ε, with four of eight of these mAbs binding to all eight strains. Although individual mAbs did not promote phagocytosis, combinations of mAbs recognising distinct epitopes either on the same domain or over different domains did. None of the mAbs inhibited IEs from binding to CSA. Our findings suggest that a combination of mAbs recognising more than one epitope would be needed for a therapeutic aiming to promote parasite clearance by phagocytosis; that DBL5ε could be considered for a VAR2CSA vaccine that aims to elicit cross-reactive antibodies that promote phagocytosis; and that identification of binding-inhibitory mAbs requires thoughtful B-cell bait design.

## Introduction

The malaria parasite *Plasmodium falciparum* causes placental infections in pregnant women. During pregnancy, the presence of the placenta selects for infected erythrocytes (IEs) that sequester in the maternal blood spaces of the placenta, triggering local inflammation and pathological remodelling of the placental tissue, disrupting normal placental function (reviewed in [1]). Malaria in pregnancy causes intrauterine growth restriction, perinatal loss, miscarriage, stillbirth, maternal anaemia and maternal morbidity in endemic areas (reviewed in [2,3])

Placental sequestration of IEs is mediated by the parasite protein VAR2CSA, a member of the cytoadhering *P. falciparum* erythrocyte membrane protein 1 (PfEMP1) family, which binds to the placental glycosaminoglycan receptor chondroitin sulfate A (CSA) [4–6]. PfEMP1 is the prominent target of antibodies to IEs [7], and antibodies to VAR2CSA have been shown to block CSA adhesion and promote parasite clearance through opsonic phagocytosis [8–11]. Women acquire protective antibodies to VAR2CSA over successive pregnancies, with multigravidae mostly having higher titres and being more protected against adverse outcomes [5,11]. Conversely, primigravidae lack VAR2CSA antibodies and are more susceptible to placental malaria and its adverse consequences. The acquisition of VAR2CSA-specific immunity with exposure and associations between VAR2CSA antibodies and protection together inspire confidence that VAR2CSA-targeted interventions could provide protection in primigravid women. Nevertheless, the large size of VAR2CSA (∼350 kDa) makes it an infeasible vaccine construct, and its high interclonal diversity [12] challenges the design of a broadly protective vaccine or therapeutic.

While vaccines based on CSA-adhering sub-domains of VAR2CSA were found to be safe, well-tolerated, and immunogenic, antibody responses have been strain-specific and failed to elicit cross-reactive immunity [13,14]. Naturally acquired VAR2CSA antibodies can be cross-reactive and inhibit CSA adhesion by heterologous parasite variants [15], suggesting conserved functional epitopes exist. The identification of these epitopes could facilitate the development of a feasible vaccine or therapeutic against VAR2CSA.

Monoclonal antibodies (mAbs) are useful tools for investigating the targets recognised by protective antibodies. Recently, two mAbs against severe malaria-associated Endothelial Protein C Receptor-binding PfEMP1 showed broad EPCR-binding inhibition activity [16], suggesting that antibodies to conserved sequestration-mediating regions of PfEMP1 can develop with natural exposure and may be leveraged to identify conserved immunogens for vaccine design. Only eight human mAbs to VAR2CSA have previously been identified, seven of which target the DBL3X and DBL5ε domains, with one mAb targeting a conserved conformational epitope spanning multiple domains [17]. To date, no adhesion-blocking VAR2CSA mAb has been identified and how VAR2CSA mAbs contribute to phagocytosis has not been investigated in detail. Here, we isolated VAR2CSA-specific B cells from Papua New Guinean multigravid women and generated immunoglobulin G subclass 1 (IgG1) mAbs to characterise their cross-reactivity across CSA-binding parasite isolates and their functional ability to inhibit CSA binding and promote opsonic phagocytosis of IEs.

## Results

### Identification of broadly cross-reactive VAR2CSA mAbs

To identify PBMC donors, plasma from Papua New Guinean multigravidae (n=53) enrolled in an intermittent preventive treatment study (Clinicaltrials.gov NCT05426434) was screened for IgG reactivity to recombinant full-length (FL) VAR2CSA by ELISA (S1 Figure). Most women (89%) had VAR2CSA-specific IgG, and PBMCs were obtained from 11 higher responders.

To identify human mAbs, B cells expressing IgG B cell receptors (BCRs) from three PBMC donors with the highest anti-VAR2CSA IgG were sorted based on dual reactivity to two fluorescently labelled FLVAR2CSA probes. Sequences of 16 cross-reactive B cells from one of the three donors (S2 Figure) yielded 16 mAbs that demonstrated dose-dependent reactivity to both FCR3 and NF54 FL-VAR2CSA proteins by ELISA (**Figure 1A**). Seven of these mAbs (named mPM1.4, 1.5, 1.7, 1.8, 1.12, 1.14 and 1.15) were cross-reactive to the two other recombinant FL-VAR2CSA tested, HB3 and Malayan Camp, while four bound just one VAR2CSA variant, and five did not bind to either the HB3 or Malayan Camp FL-VAR2CSA. Fourteen of the 16 new mAbs bound to FL-VAR2CSA of FCR3 and NF54 at considerably lower concentrations than the previously published mAb PAM2.8 [17] (**Figure 1A**). When mAbs were tested for binding to VAR2CSA sub-domains, eight mAbs targeted the DBL3X domain, and the other eight targeted the DBL5ε domain of VAR2CSA, with none binding DBL1X-DBL2X-ID2a, DBL4ε or DBL6ε (**Figure 1B**).

**Figure 1.**
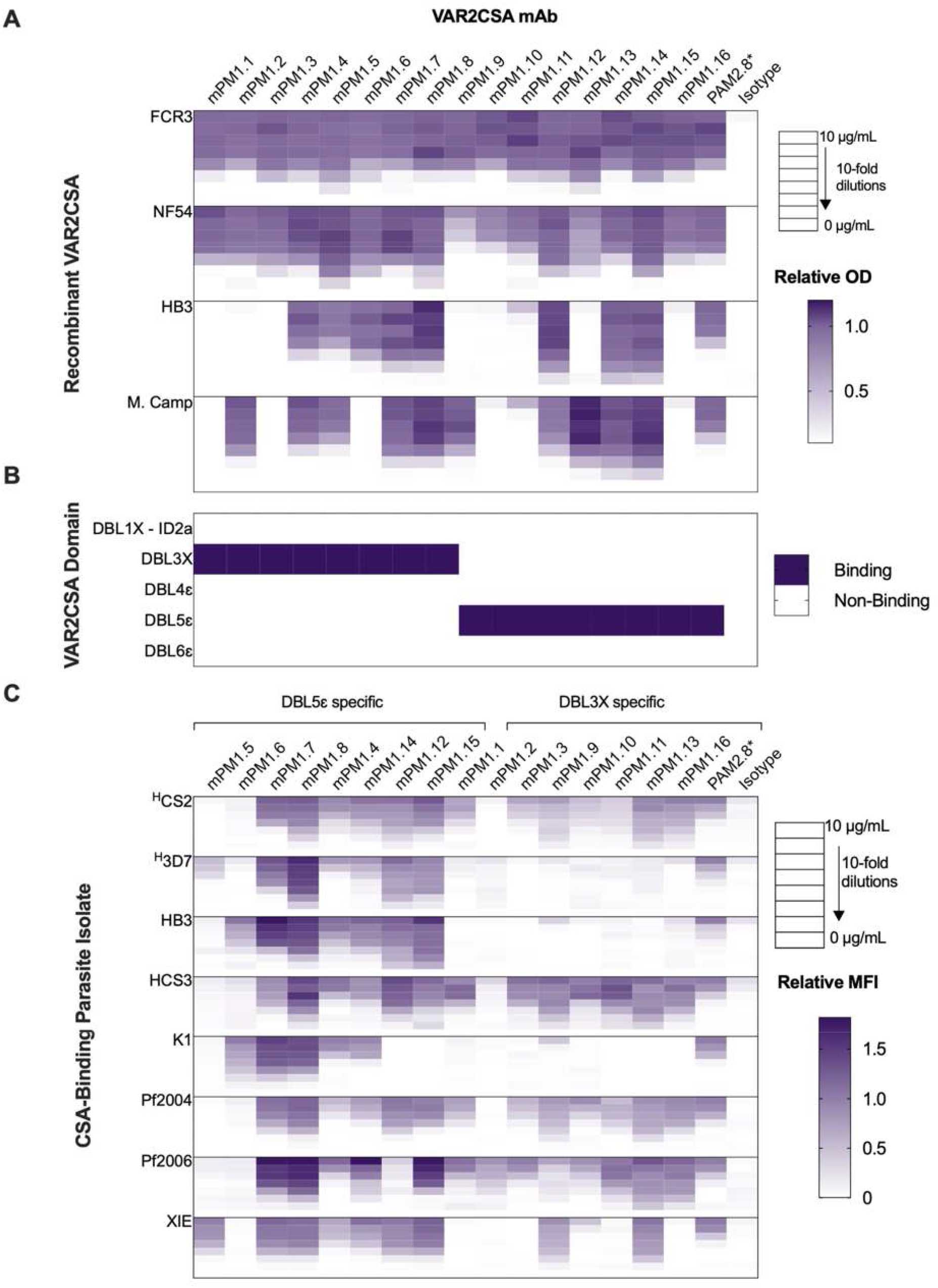
Binding Profiles of VAR2CSA mAbs. (A) Heatmap of mAb binding to full-length VAR2CSA proteins in a dose-dependent manner. Full length VAR2CSA proteins from FCR3 and NF54 strains (used to isolate antigen-specific B cells) and VAR2CSA of HB3 and Malayan Camp strains were used in ELISAs. mAb binding is represented as optical density (OD) relative to PAM2.8 control mAb at 10 µg/mL. (B) Heatmap of mAb binding to VAR2CSA DBL3 and DBL5 domain constructs determined by the area under the dose-dependent curve (AUC), with binding defined as AUC > 4 based on the isotype control. (C) Heatmap of mAb recognition of native VAR2CSA on heterologous CSA-binding isolates in a dose-dependent manner. mAb recognition measured as mean fluorescence intensity (MFI) relative to PAM2.8 at 10 µg/mL. *Previously published VAR2CSA mAb. ^H^Isolates expressing homologous VAR2CSA to FCR3 (subclone CS2) and NF54 (subclone 3D7) variants that were used to isolate VAR2CSA-specific B cells.

Glycan profiling of four of the expressed mAbs (mPM 1.13, 1.14, 1.15, 1.16) by mass spectrometry shows that almost all complex glycans of these mAbs contained a core fucose (S1 Table).

To assess whether the mAbs recognise native VAR2CSA on the surface of IEs and whether they are cross-reactive with different CSA-binding parasite isolates, we used eight CSA-binding laboratory isolates. Each was confirmed to be distinct through genotyping of *glurp* and *msp2* merozoite genes (S3 Figure), and their CSA-binding phenotype was enriched and verified (S4 Figure). Flow cytometry analysis of mAb binding to these IEs confirmed that all mAbs recognised native VAR2CSA on IEs, although recognition by individual mAb varied with each CSA-binding isolate. Individual mAb reactivity to parasite isolates expressing the homologous native protein, CS2 (a subclone of FCR3) and 3D7 (a subclone of NF54), varied between mAbs, with some recognising one homologous native variant more than the other. Two mAbs, mPM1.6 and mPM1.2, recognised heterologous better than homologous VAR2CSA on IE (**Figure 1C**), and four of the novel DBL5ε-specific mAbs (mPM1.7, 1.8, 1.4, 1.14) were broadly cross-reactive across all eight tested CSA-binding parasite isolates (**Figure 1C**). Most of the novel mAbs bound to IEs at lower concentrations than PAM2.8 did. As expected, a non-CSA binding parental line of CS2, E8B [18], was not recognised by any of the seven mAbs tested (S5 Figure).

### Broadly reactive mPM1.7 targets a conformational epitope

To identify the potential epitope of mPM1.7 (which recognises all tested VAR2CSA variants), hydrogen-deuterium exchange mass spectrometry (HDX-MS) was utilised, and residues with significant changes in solvent exchange in the presence of the mAb (S8 Figure) were mapped onto the previously published cryo-EM structure of DBL5ε-DBL6ε region (NF54) [19]. Notably, several residues had a decrease in HDX in the presence of the antibody and were located in two regions of the protein. The first region, ‘R1’, consisted of residues L146-N152 and the second region, ‘R2’, consisted of residues K52, Y122, M129-N132, Q134-F135 and L200-E205, where all of these residues come together to form a cavity (**Figure 2A**). These same residues were also highly conserved across 17 sequences of DBL5ε domains [20](**Figure 2B**).

**Figure 2.**
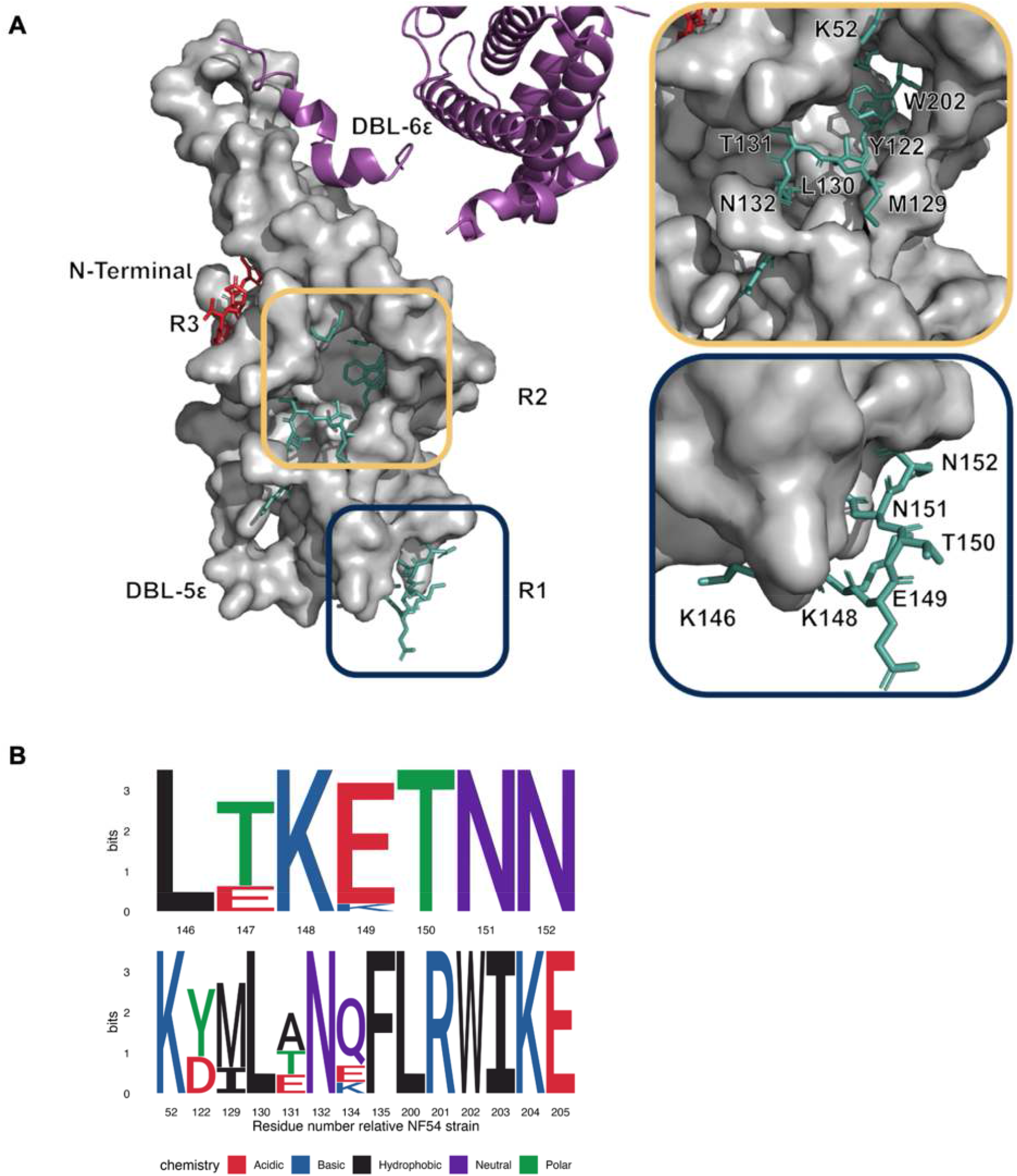
DBL5ε peptides with changes in deuterium exchange upon complex formation with mPM1.7. (A) Residues of peptides with decreased (R1 and R2) and increased (R3) deuterium exchange mapped onto the published cryo-EM structure of DBL5ε-DBL6ε (PDB:7JGG). (B) Sequence logo plot showing amino acid conservation of DBL5ε peptides with decreased deuterium exchange, R1 (top) and R2 (bottom), with PAM1.7 complex formation across 17 published sequences, coloured by chemistry and numbered relative to the NF54 DBL5ε sequence. The R2 (bottom) plot displays non-consecutive residues of the primary sequence that are adjacent in the tertiary structure of the protein.

### mAbs did not inhibit adhesion of IEs to CSA

The ability of the mAbs to inhibit adhesion of IEs to CSA was tested. CS2-IEs were opsonised with each individual mAb or with a pool of all 17 mAbs (including PAM2.8) in combination. IEs opsonised with pooled immune plasma or soluble CSA were inhibited from binding CSA at 90% and 75% inhibition, respectively (**Figure 3**). Neither individual mAbs nor a combination of all mAbs inhibited IE: CSA adhesion.

**Figure 3.**
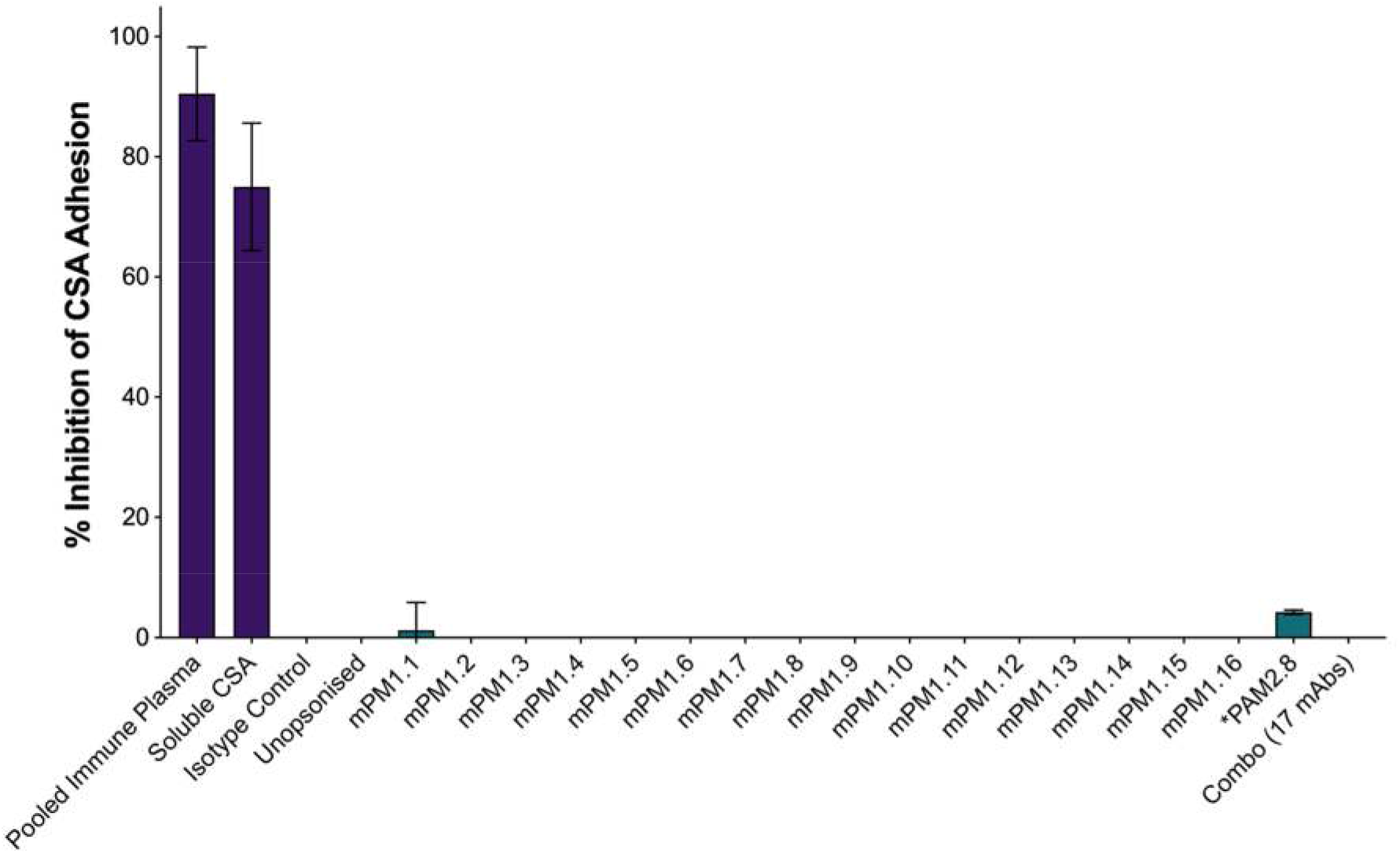
VAR2CSA-specific mAb-mediated inhibition of CSA binding by placental-type IEs. Inhibition of CS2-IE binding to CSA under static conditions by individual mAbs and all mAbs in combination (combo) at a total mAb concentration of 10 µg/mL, measured by colourimetry. *Previously published VAR2CSA mAb [17]. Data are mean ± SD from two experiments.

### mAbs targeting distinct regions synergistically enhance phagocytosis of IEs

The ability of mAbs to promote phagocytosis of IEs by THP-1 cells was also assessed. When CS2 IEs were opsonised with 10 µg/mL of any single mAb, there was no phagocytosis above that of the isotype control (**Figure 4**). However, a pool of all 16 new mAbs plus PAM2.8 at a total mAb concentration of 10 µg/mL (each mAb at 0.6 µg/mL) showed 35% phagocytosis, while the pooled immune plasma resulted in 50% phagocytosis of CS2-IEs (**Figure 4**).

**Figure 4.**
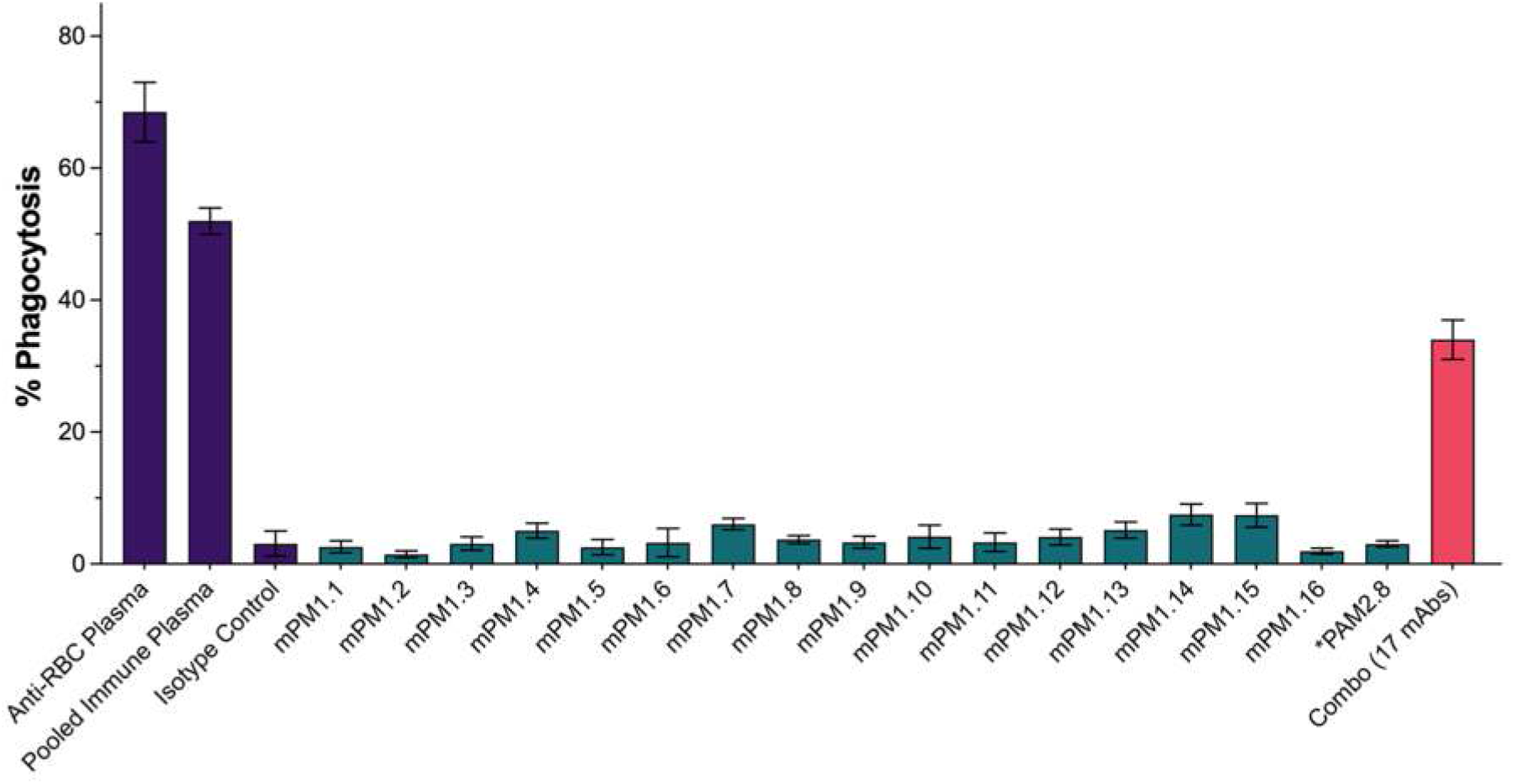
VAR2CSA-specific mAb-mediated phagocytosis of CSA-binding IEs by THP-1 cells. Phagocytosis of CS2-IEs opsonised with individual mAbs (teal) or all mAbs in combination (pink) at a total mAb concentration of 10 µg/mL was measured using flow cytometry. *Previously published VAR2CSA mAb. Anti-RBC Plasma: Rabbit plasma against human red blood cells. Pooled Immune Plasma: Plasma pool with high recognition of CS2-IEs. Isotype Control: IgG1 mAb against influenza HA protein. Bars indicate mean ± SEM from two experiments.

Given the pool of mAbs and pooled immune plasma both induced phagocytosis and contained multiple epitope specificities, this suggested that engaging multiple epitopes on VAR2CSA simultaneously with mAbs is likely required for efficient phagocytosis. Therefore, we sought to determine the minimal set of mAbs that could simultaneously bind VAR2CSA to maximally opsonise the protein. To map epitope overlap among domain-specific mAbs, competition ELISAs were conducted separately for DBL3X and DBL5ε-specific mAbs. Competition (inhibition of antigen binding) of all tested pairwise combinations (S6 Figure) are displayed in competition networks (**Figure 5**). Three of eight DBL3X-specific mAbs, mPM1.2, 1.9, and 1.13, competed with each other but did not compete with the five other mAbs, mPM1.1, 1.3, 1.10, 1.11, and 1.16 (**Figure 5A**). Consistent with this, mPM1.1, 1.3, 1.10, 1.11, 1.16 and PAM2.8 competed with each other but did not compete with mPM1.2, 1.9, and 1.13 (**Figure 5A**). This indicates that mPM1.2, 1.9, and 1.13 target a distinct region of DBL3X to mPM1.1, 1.3, 1.10, 1.11, 1.16, and PAM2.8. Out of the eight DBL5ε-specific mAbs, mPM1.6, 1.7, and 1.8 competed with each other and with the other five DBL5ε mAbs (**Figure 5B**). mPM1.5, 1.12, and 1.15 competed with each other and with mPM1.6, 1.7, and 1.8; however, they did not compete with mPM1.4 or 1.14. (**Figure 5B**). This suggests these mAbs target two overlapping regions of DBL5ε, with mPM1.5, 1.12, and 1.15 targeting a different region from that of mPM1.4 or 1.14, and mPM1.6, 1.7, and 1.8 targeting a region overlapping with the targets of mPM1.5, 1.12, 1.15, 1.4, 1.4, or 1.14.

**Figure 5.**
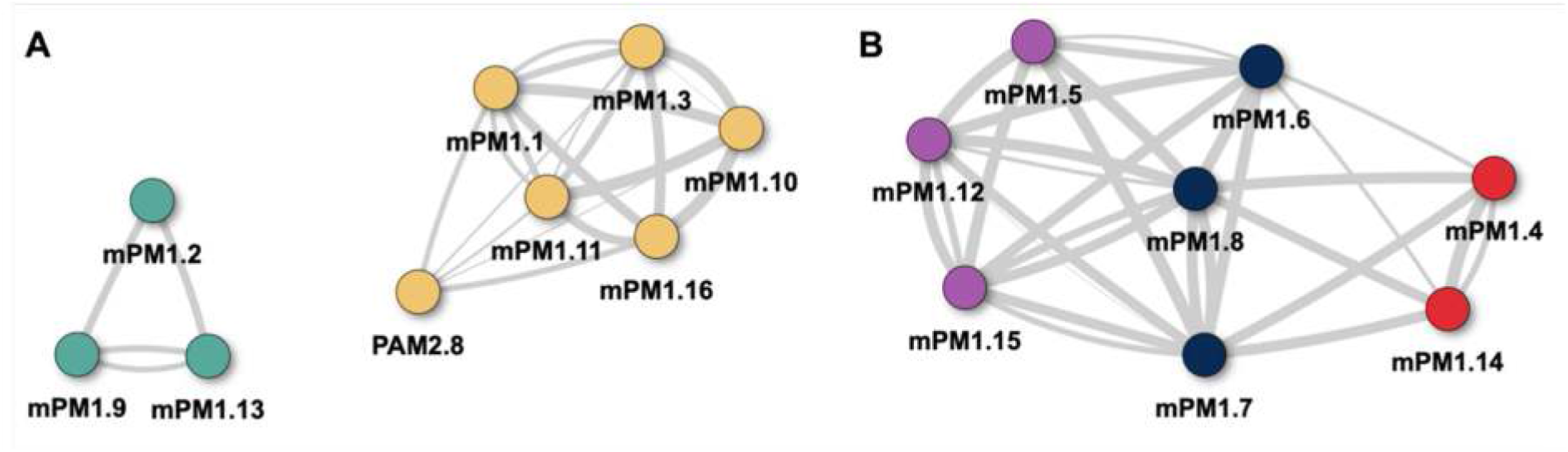
Competition between mAbs targeting the same VAR2CSA domain. Competition networks of (A) DBL3X-specific and (B) DBL5ε-specific mAbs illustrate how mAbs inhibit each other’s antigen binding. mAb clustering is based on competition ELISAs, with competition defined as >50% inhibition of antigen binding, shown by connecting lines between two mAbs, where line thickness indicates the strength of competition. mAbs sharing the same colour compete with the same antibodies.

Non-competing mAbs targeting VAR2CSA were combined into cocktails to assess their effects on inducing phagocytosis of CS2-IEs by THP-1 cells. Four different cocktails were created of three or four mAbs.

Cocktails of three or four non-competing mAbs enhanced phagocytosis of IEs to levels similar to the combination of all 17 mAbs (**Figure 6A**). Compared to individual mAbs, the levels of phagocytosis induced by the cocktails were greater than the combined activity of the individual mAbs making up each cocktail, indicating synergy between mAbs. The amount of IgG on the surface of CS2-IEs when opsonised by the mAb cocktails of three or four non-competing mAbs were also comparable to those of the 17 mAb combination (S7 Figure). Opsonisation cocktails were compared over multiple opsonisation dilutions, with the cocktails containing four mAbs showing slightly higher phagocytosis over multiple dilutions compared to cocktails containing three mAbs (**Figure 6B**).

**Figure 6.**
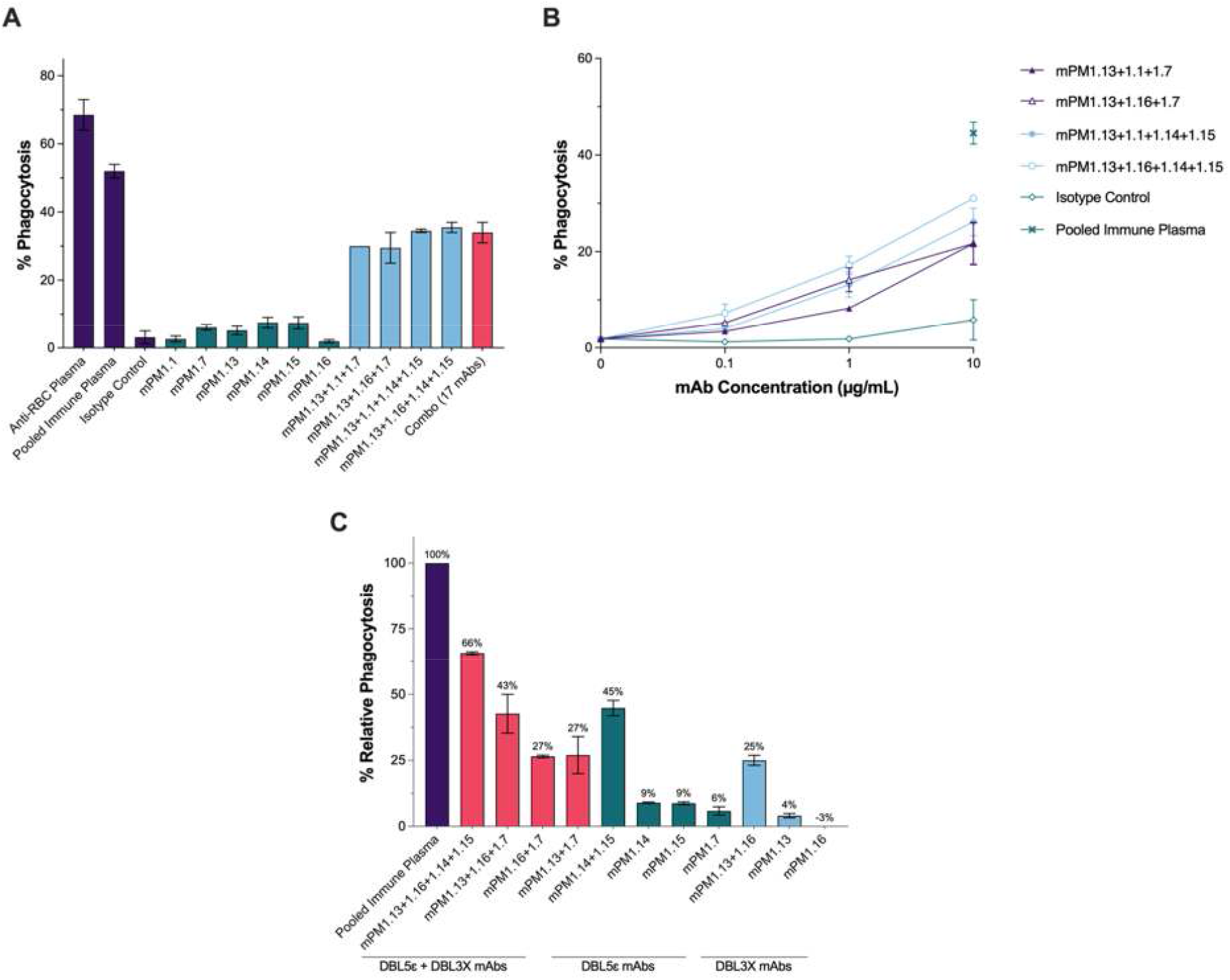
Characterisation of VAR2CSA mAb-mediated phagocytosis of IEs by THP-1 cells. (A) Phagocytosis of CS2-IEs opsonised with a single mAb (teal), a cocktail of non-competitive mAbs (blue), or a combination of 17 mAbs, including PAM2.8 (pink), at a total mAb concentration of 10 µg/mL. (B) Dose-dependent phagocytosis of CS2-IEs opsonised with cocktails of non-competitive mAbs at total mAb concentrations of 10, 1 and 0.1 µg/mL. (C) Phagocytosis of CS2-IEs opsonised with 10 µg/mL of mAbs specific to DBL3X alone (blue), DBL5ε alone (teal), both DBL3X and DBL5ε (pink), and cocktail including and excluding broadly reactive mPM1.7 normalised to phagocytosis relative to pooled immune plasma. Mean ± SEM from two experiments

How monoclonal combinations to single or multiple domains could promote phagocytosis was also investigated. A cocktail of DBL3X-specific mAbs mPM1.13 and mPM1.16 or DBL5ε-specific mAbs mPM1.15 and mPM1.14 resulted in phagocytosis of 25% and 45%, respectively. When opsonising with a cocktail containing these four mAbs, phagocytosis increased further (66%) (**Figure 6C**). Similarly, combining a single DBL3X mAb (mPM1.13 or mPM1.16) with a single DBL5ε mAb (mPM1.7) increased phagocytosis compared to each mAb alone (**Figure 6C**).

## Discussion

The VAR2CSA PfEMP1, which mediates IE sequestration in the placenta, is the main target of antibodies that protect against placental infection and adverse outcomes during *P. falciparum* infections in pregnancy. Whilst naturally acquired functional antibodies to VAR2CSA correlate with protection, vaccine development efforts have been hampered by antigenic diversity among isolates, and clinical trials of vaccines based on VAR2CSA subunits have shown that vaccination-induced antibodies do not show evidence of recognising VAR2CSA on heterologous IEs [13,14]. Although anti-VAR2CSA mAbs have been previously isolated, these failed to inhibit CSA adhesion. However, naturally acquired polyclonal antibodies can broadly inhibit CSA adhesion (Doritchamou et al., 2022), suggesting that such key protective mAbs could exist. Here, we show that multigravidae develop mAbs capable of broad recognition and synergistic opsonic phagocytosis; all identified mAbs targeted immunodominant domains that do not directly participate in CSA adhesion [19], and none of them inhibited CSA binding.

A total of 16 novel mAbs were identified in this study from a single multigravid donor in PNG, with half targeting the DBL3X domain and the other half targeting the DBL5ε domain. Previous efforts by Barfod *et al*. to isolate mAbs against FL VAR2CSA from a Ghanaian cohort yielded similar findings: seven of the eight mAbs, one of which was PAM2.8, targeted either the DBL3X or DBL5ε domains, and one mAb, PAM1.4, targeted a conformational epitope spanning domains ID1, DBL2X, ID2, and DBL4ε [17,21]. These findings are consistent with DBL3X and DBL5ε being immunodominant as malaria-exposed multigravidae develop IgG predominantly against DBL3X and DBL5ε [22,23]. In addition, animal immunisation studies with these domains show that antibodies towards them are strongly cross-reactive compared to other VAR2CSA domains [24], and it is possible that our strategy of using two heterologous baits biased our selection towards B cells that recognise these two domains.

All 16 mAbs recognised native VAR2CSA expressed on IEs, but strong reactivity to recombinant protein did not necessarily predict reactivity to the native homologous variant and vice versa. For example, mPM1.5 exhibited dose-dependent reactivity to the FL HB3 recombinant protein but failed to recognise the HB3 isolate. Likewise, mAbs mPM1.6 and mPM1.2 isolated using FCR3 and NF54 variant baits did not show dose-dependent reactivity against the homologous isolates CS2 and 3D7 (expressing FCR3 and NF54-like VAR2CSA), but did recognise VAR2CSA on other isolates (HB3, K1, Pf2006). This may be due to epitopes of the recombinant proteins being conformationally different from those on the parasite surface due to the expression systems used, or potentially the sequences of the isolates in our lab are different from those of the reference strain sequences used to make the recombinant proteins.

In general, the mAbs we identified to DBL5ε were more cross-reactive compared to mAbs to DBL3X: all four of the mAbs which recognised all isolates targeted DBL5ε, and the only DBL3X mAb that was broadly reactive against all isolates was the previously described PAM2.8, which we used as a control [17,25,26]. The highly potent, broadly reactive mAb, mPM1.8, was clonally related to mPM1.7, sharing similar heavy and light chain immunoglobulin gene usage, nearly identical heavy chain complementary determining region 3 (CDR-H3) sequences, and a light chain CDR-L3 that differed by only 3 amino acids (S2 Table). Hence, the more potent of the two, mPM1.8, may have resulted from an affinity-matured B cell within the shared clonal lineage. The superior cross-reactivity of DBL5ε-specific mAbs compared to DBL3X-specific mAbs is partly explained by higher DBL5ε sequence conservation. In previous studies, geographically distinct isolates had an average amino acid identity of 87% (range of 83-99%) for DBL5 [20], and DBL5ε is the second most conserved of the VAR2CSA domains after DBL4ε [27].

Epitope mapping of the broadly reactive mAb mPM1.7 using HDX-MS revealed two regions of DBL5ε protected from deuterium exchange upon mPM1.7 binding. The first region (R1, **Figure *2*A**) is well-exposed to the solvent and sterically accessible; thus, a decrease in deuterium exchange is consistent with binding. The second region (R2, **Figure *2*A**) forms a cavity in the folded protein. Whilst some residues are near the “face” of the protein, several are sterically inaccessible. Therefore, it is possible that a few residues near the face of this cavity directly interact with mPM1.7, while several others within the cavity are blocked from solvent access upon mPM1.7 binding. Although HDX data does not distinguish between true epitope-defining residues and auxiliary residues, and our data have peptide-level resolution rather than single-amino-acid resolution, our finding is consistent with mPM1.7 targeting two distinct regions of the folded protein, suggesting that the epitope is conformational rather than linear. An additional third region, consisting of residues W39-D43 (R3, **Figure *2*A**), showed greater exchange in the complex than in the uncomplexed analyte. This is indicative of a slight conformational change to the protein that increased solvent accessibility for this region.

Competition for antigen binding among the novel mAbs indicates that our DBL3X and DBL5ε mAbs recognise at least two distinct regions within their respective domains, suggesting that at least two novel mAbs can target a single VAR2CSA domain simultaneously. The competition patterns of DBL5ε-reactive mAbs also indicated that some DBL5ε mAbs were targeting an overlapping region between those distinct regions on DBL5ε. Previous VAR2CSA DBL3X mAbs were also shown to recognise antigenically distinct epitopes, whereas DBL5ε mAbs have been reported to target only overlapping epitopes [17].

Opsonic phagocytosis of IEs by THP-1 cells was negligible in the presence of a single opsonising mAb. Even the PAM2.8 mAb previously reported to promote THP-1 phagocytosis of FCR3 (29.1%) and HB3 IEs (∼10%) [25] did not promote phagocytosis of CS2 IEs in our hands. However, combining as few as two non-competing mAbs at matched antibody concentrations synergistically increased IE uptake by THP-1 cells, with overall phagocytosis levels further improving with the addition of a third or fourth mAb. The reason for this inconsistency with PAM2.8 is not fully understood, especially given that FCR3 and CS2 isolates are closely related (Avril et al., 2010; Duffy et al., 2005). PfEMP1s are structurally anchored within protrusions known as knobs on the IE surface [29]. It is possible that the CS2 isolate may have a sparser distribution of VAR2CSA per given knob than the FCR3 isolate used by Barfod *et al*. [29]. We hypothesise that the VAR2CSA density on the surface of IEs is a limiting factor for an antibody’s ability to effectively engage and cross-link as required for FcR-mediated phagocytosis, as has been described for complement fixation on IEs [30], and that the presence of multiple mAb/PfEMP1 is needed to promote IE phagocytosis.

The mAbs we describe could promote phagocytosis by targeting either DBL3X or DBL5ε alone or in combination. This is consistent with previous work showing that IgG from malaria-exposed multigravidae with strong phagocytic activity against CSA-binding IEs predominantly target DBL3X and DBL5ε (Lambert et al., 2014).

The cross-reactive nature of the DBL5ε mAbs, which also promote phagocytosis of IEs is noteworthy, as it provides further evidence that antibodies to the DBL5ε domain can provide functional cross-reactive immunity [22,24]. We have previously shown that there are multiple pathways to protection, and the ability of antibodies to promote phagocytosis is associated with a reduced risk of placental malaria [8]. Though it may be ideal to have a vaccine or monoclonal therapeutic that can both inhibit parasite binding to the placenta and promote IE clearance by phagocytosis, DBL5ε could be considered for inclusion in VAR2CSA vaccines due to its potential to induce cross-reactive and functional IgG.

None of the mAbs we isolated were able to inhibit binding of the parasites to the placental receptor CSA. This may be because they bound to either DBL3X or DBL5ε and are therefore unlikely to sterically interfere with CSA binding, as the domains associated with the CSA binding groove of VAR2CSA are DBL1, DBL2 and DBL4 [19,31]. Likewise, previously published mAbs do not inhibit CSA binding, nor recognise regions associated with the CSA binding groove [25].

We chose to use FL VAR2CSA as baits as cross-reactive antibodies are known to recognise structural epitopes available on the quaternary structure [15]. Though this strategy produced cross-reactive antibodies, it did not yield antibodies with the desired CSA blocking activity. Future efforts to identify binding inhibitory B cells should focus on identifying mAb targeting DBL1X, DBL2X and DBL4ε domains, while avoiding the immunodominant DBL3X and DBL5ε, for example, by using recombinant DBL3X and DBL5ε as negative baits.

In summary, we describe 16 new mAbs to VAR2CSA. Our data suggests that clearance of IEs via Fc-mediated effector functions, such as opsonic phagocytosis, depends on the combined activity of antibodies targeting multiple epitopes (either on the same domain or across multiple domains) rather than individual monoclonals. This has important implications for therapeutic strategies, indicating that combinations of mAbs may be required to achieve functional efficacy. For vaccine design, our data support the inclusion of broadly conserved VAR2CSA epitopes such as those on DBL5ε that elicit strong Fc-dependent effector functions, in addition to those that inhibit CSA adhesion.

## Materials and Methods

### Study Participants

Participants were a subset of women enrolled in the study, “SAPOT” (Clinicaltrials.gov NCT05426434), which ran from 2022 to 2025. Enrolment in the sub-study and PBMC collection occurred in May-June 2023. All participants provided written informed consent. The research ethics for this sub-study were approved by the Institute of Medical Research - Institutional Review Board (IRB# 2301), the PNG Medical Research Advisory Committee (MRAC# 22.69) and the University of Melbourne Office of Research Ethics and Integrity (Ref# 2023-25683-41954-3).

Plasma samples of 53 multigravidae taken at delivery in the SAPOT study from Madang, PNG, were screened using ELISA [32] to identify participants with VAR2CSA IgG. Eleven seropositive women were recruited for this study, and whole-blood samples were collected >6 weeks post-partum for PBMC isolation. Three PBMC samples from individuals with the highest IgG levels to FL VAR2CSA by ELISA were selected to screen for VAR2CSA-specific B cells.

### Recombinant Proteins

Four variants of FL VAR2CSA, five VAR2CSA domain constructs, PAM2.8 and IgG1 isotype control antibodies were used in experiments (S3 Table). Recombinant VAR2CSA for strains FCR3 and NF54 were prepared in-house. Codon-optimised expression constructs were synthesised with an N-terminal bovine prolactin signal peptide and a C-terminal His tag (GSHHHHHH) (GeneArt, ThermoFisher), and cloned into mammalian expression vectors. VAR2CSA proteins were expressed by transient transfection of Expi293 cells using ExpiFectamine (Thermo Fisher) and purified from culture supernatant using Ni Sepharose Excel resin and size exclusion chromatography in PBS on a Superdex 200 pg 16/600 column (Cytiva).

### Isolation of cross-reactive VAR2CSA B-cells

VAR2CSA-specific B cells were identified within cryopreserved PBMC samples by co-staining with fluorescently labelled FCR3 and NF54 strain VAR2CSA proteins prepared using Lightning-Link PE and APC conjugation kits (Abcam), respectively. The staining panel included IgG BV786 (G18-145, dilution 1:50), IgD PE-Cy7 (IA6-2, dilution 1:333) (BD), CD19 ECD (J3-119, dilution 1:100, Beckman Coulter, Brea, USA) along with BV510 dump makers (CD14, M5E2, dilution 1:200; CD3, OKT3, dilution 1:400; CD8α, RPA-T8, dilution 1:1000; CD16, 3G8, dilution 1:333; CD10, HI10a, dilution 1:500; all from BioLegend). Cell viability was determined using Aqua Live/Dead amine-reactive dye (Thermo Fisher). B cells were single cell-sorted into 96-well PCR plates and immediately frozen on dry ice and stored at -80°C until sequencing.

### Sequencing, cloning and expression of recombinant IgG

The sequencing and cloning of BCRs from single B cells was performed as previously described [33,34]. Plasmids expressing heavy and light immunoglobulin chains were transfected into Expi293F cells using ExpiFectamine (Thermo Fisher). Recombinant monoclonal antibodies were purified from culture supernatants using MabSelect PrismA protein A resin (Cytiva) as per the manufacturer’s instructions, and then buffer exchanged into PBS. Afucosylation of mAb were determined using mass-spec (See S1 Methods)

### Parasite culture, selection and validation of CSA-binding phenotype

*P. falciparum* isolates were cultured in medium containing 5% human serum and 0.25% Albumax II and selected for CSA binding phenotype as previously described [35](S2 Methods). A published colourimetric binding inhibition assay was adapted to measure CSA binding levels in enriched parasite isolates [9].

Parasite isolates Pf2004, Pf2006, and K1, which initially showed limited CSA binding, were panned on BeWo cells to enrich for CSA-binding parasites before selection was continued with immobilised CSA. The selection process was repeated three times to obtain a higher number of CSA-binding parasites.

### Genotyping of *P. falciparum* isolates

Unique lab isolates were confirmed by investigating polymorphic regions of msp2 and glurp by nested PCR using primers and amplification conditions as described by Ullah and colleagues [36]. Parasite DNA from each isolate was extracted using the QIAamp DNA blood Kit (QIAGEN) according to the manufacturer’s protocol.

### mAb reactivity to variants of native VAR2CSA on IEs

Flow cytometry was used to evaluate mAb recognition of native VAR2CSA on the surface of IEs [37]. Briefly, trophozoite-stage IEs enriched by gelatin floatation [38] at 0.2% haematocrit were opsonised with 10-fold serial dilutions of mAbs from 10 µg/mL in 0.1% BSA in PBS (assay and wash buffer) for 1hr at RT before being washed and stained with 64 µg/mL of polyclonal rabbit anti-human IgG (Dako, Agilent) for 30 mins at RT. Cells were washed again and stained with 2 µg/mL polyclonal donkey anti-rabbit IgG-Alexa Fluor 647 (Life Technologies) and 1X SYBR Green (Invitrogen) for 30 mins at RT, then washed and finally fixed with 2% PFA and 0.075% glutaraldehyde for at least 15 mins prior to analysis by flow cytometry.

### mAb functional assays

mAb-dependent inhibition of IE binding to immobilised CSA [9]. One million Percoll-purified IEs were opsonised with 10 µg/mL of an individual mAb or a combination of mAbs and transferred to CSA-coated wells in duplicate. Opsonisation with pooled immune plasma and soluble CSA for assay controls was performed at 1:10 dilution and 100 µg/mL, respectively.

THP-1 phagocytosis of mAb-opsonised CS2 IEs was assessed using flow cytometry as previously described [39].

### Measurement of competition between mAbs

To assess whether mAbs targeting the same domain of VAR2CSA shared overlapping epitopes, a competition ELISA was performed. Individual mAbs were biotinylated using the EZ-Link Sulfo-NHS-LC-Biotin kit (Thermo Fisher). ELISAs were performed as previously described [32], but with the addition of a competitor mAb. VAR2CSA (0.5 ug/mL) was coated onto a 96-well flat-bottom plate. 0.1 µg/mL of biotinylated mAb and 5 µg/mL of competitor were pre-mixed before adding onto antigen-coated wells and incubated for 1hr at RT. The binding of mAb-biotin was detected with Streptavidin-HRP and TMB substrate. mAbs targeting the same domain were tested in reciprocal assays, with each acting as both detector and competitor.

### HDX-MS analysis of mAb: VAR2CSA interactions

The DBL5ε domain (3D7) was analysed by HDX-MS in the absence and presence of equimolar mPM1.7 using a PAL Dual Head HDX Automation manager (Trajan/LEAP) controlled by ChronosHDX software (Trajan). Samples (4 µL) were incubated in 55 μL of either a deuterated (50 mM potassium phosphate Buffer pD 7.0, 150 mM NaCl in D_2_O) or non-deuterated phosphate buffer (50 mM potassium phosphate Buffer pH 7.4, 150 mM NaCl in H_2_O). Reactions were quenched by adding 50 μL of the protein to 55 μL of quench buffer (50 mM potassium phosphate buffer, pH 2.3, containing 4M guanidine hydrochloride and 200 mM Tris(2-carboxyethyl)phosphine hydrochloride). Online digestion was then performed by passing 95 µL of the quenched sample through an immobilised pepsin column (2.1 m x 30 mm, Waters) equilibrated in 0.1 % formic acid at 100 μL/min. Peptides were captured and desalted by a C18 trap column (VanGuard BEH; 1.7 μm; 2.1 × 5 mm(Waters)) before elution and separation on an ACQUITY UPLC BEH C18 analytical column (1.7 μm, 1 × 100 mm (Waters)). Mass spectrometry was performed on a Synapt G2-Si (Waters). Instrument settings were: 3.0 KV capillary and 40 V sampling cone with source and desolvation temperatures of 100 and 40 °C, respectively. The desolvation and cone gas flow were at 80 L/hr and 100 L/hr, respectively. All mass spectra were acquired using a 0.4 s scan time with continuous lock mass (Leu-Enk, 556.2771 *m/z*) for mass accuracy correction. Data were acquired in MS^E^ mode with a ramp from 20 to 40 V. The DynamX software was used to analyse the results, and additional visualisation was performed in PyMOL [40].

### Data analysis

mAb binding and functional data were processed in Microsoft Excel and visualised using GraphPad Prism 10. For competition ELISA data, % inhibition of antigen binding was calculated relative to the self-competitor (the detected biotinylated mAb and the competitor mAb were the same). Clustering networks based on >50% competition/inhibition were generated and visualised using the ggraph and visNetwork packages in R, respectively. Sequence logo plots for the conserved DBL5ε residues were prepared from aligned sequences using the ggseqlogo package and visualised with the ggplot2 package in R.

## Acknowledgements

This work was funded by the National Health and Medical Research Council (NHMRC) Australia GNT 2020606 awarded to SR and EA and Australian Centre Research Excellence Malaria Elimination (ACREME) seed grant awarded to EA. This was a sub-study of the SAPOT trial led by HWU and supported by NHMRC (GNT2000780). CG and AW are supported by a Cumming Global Centre for Pandemic Therapeutics grant and an NHMRC Investigator grant. We would like to thank Patrick Duffy and David Narum from NIH and NIAID, United States, and Morten Neilson, Lars Hviid, and Maria del Pilar Quintana Varon from the University of Copenhagen, Denmark, for gifting the proteins used in our assays. We would also like to acknowledge Magdaline Sakkas from the Melbourne Cytometry Platform, Brain Centre Flow Cytometry node, for help with flow cytometry, and the Bio21 Mass Spectrometry and Proteomics Facility for help with HDX-MS. Thank you to Phantica Yambo from the PNGIMR and Olivia Wilhelm from PDI for technical assistance, and Grant Lee from Menzies for help with data. Finally, thank you to the women in Madang, Papua New Guinea, who generously gave both their time and PBMC to enable this work to go ahead.

## Supporting Information Captions

**S1 Table : % afucosylation for mAbs S2 Table: VAR2CSA mAb Sequences**

**S3 Table: Summary of recombinant VAR2CSA proteins and mAb controls**

**S1 Figure: VAR2CSA antibody screen of participants of the SAPOT parent study**. Plasma samples from participants in the SAPOT study (n=53) were screened by ELISA to identify multigravidae with high VAR2CSA antibody levels. A subset of seropositive women (n=11) was recruited with informed consent, and their peripheral blood mononuclear cells (PBMCs) were collected for the present study. Three of the donors with the highest IgG levels to VAR2CSA were screened for B cells that cross-reacted with the NF54 and FCR3 VAR2CSA antigen baits to identify mAbs.

**S2 Figure: FACS gating and VAR2CSA probe staining with annotated B cells yielding successful mAbs**. (A) Gating used to identify IgG+ B cells from PBMCs. (B, C) VAR2CSA FCR3 and NF54 probe staining on IgG+ B cells. Sorted cells that yielded sequences for mPM1.1–mPM1.16 are indicated by colour according to the legends.

The overall gate used to sort probe-positive cells is shown in pink.

**S3 Figure: Genotyping of *glurp* and *msp2* of laboratory parasite isolates**. Genotyping of parasite isolates using primers specific to (A) *glurp, (B, C) msp2* of 3D7/IC allelic family and (D) *msp2* of FC27 allelic family demonstrated that all isolates were genotypically distinct. (C) A repeat gel for P2004 products from gel (B). Parasite isolates Palo Alto, W2MEF, and TM284 were not used for mAb characterisation, as Palo Alto could not be maintained in culture and W2MEF and TM284 did not express a CSA-binding phenotype. NT: no template, bp: base pair

**S4 Figure: CSA-binding of heterologous parasite isolates**. Parasite isolates followed by -CSA were panned on immobilised CSA and/or BeWo cells to select for parasite phenotype. CSA binding of isolates to immobilised CSA was then measured as described by Dube *et al*. (2025), and binding levels were calculated relative to the CS2 isolate with constitutive expression of the CSA-binding phenotype. Bars represent mean ± SD from two individual experiments, except ✧, bars represent duplicates from an individual experiment.

**S5 Figure: Representative mAb reactivity to infected erythrocytes (IEs) of CSA-binding and non-CSA-binding parasite isolates**. mAb reactivity to (A) CS2 parasite isolate with CSA binding phenotype and (B) E8B parasite isolate that does not predominantly bind CSA. Mean Fluorescence Intensity (MFI) is adjusted by subtracting the MFI of uninfected erythrocytes from that of IEs. Indicates mean and SD from experimental duplicates. PIP: Pooled Immune Plasma

**S6 Figure: Competition matrices of mAbs targeting a single VAR2CSA domain**. The binding of a biotinylated test mAb was detected in the presence of an excess of competitor to assess whether two mAbs compete for a shared binding site. (A) Competition of mAbs targeting the DBL3X domain of VAR2CSA. (B) Competition of mAbs targeting the DBL5ε domain of VAR2CSA. Competition is calculated as optical density relative (rOD) to the self-competitor and is defined as strong (rOD ≥ 0.75), moderate (0.5 ≤ rOD < 0.75), weak (0.25 ≤ rOD < 0.5), or no competition (rOD ≤ 0.25).

*PAM2.8 previously published mAb.

**S7 Figure: mAb cocktail reactivity to CS2 IEs**. The binding of mAb cocktail formulations to IEs was evaluated to assess whether opsonisation levels varied significantly across formulations. All conditions tested were at a total antibody concentration of 10 µg/mL. Mean fluorescence intensity (MFI) is calculated relative to PAM2.8 (previously published VAR2CSA mAb) at 10 µg/mL. The graph presents the mean ± SD from two experiments.

**S8 Figure: Uptake plots for selected peptides in the three noted regions of DBL5ε**. Peptides in regions 1 and 2 demonstrate significantly decreased deuterium uptake in the presence of mPM1.7, especially at long exposure, whilst region 3 demonstrates the inverse.

**S1 Methods: Mass spectrometric determination of afucosylation of mAbs**

**S2 Methods: Parasite culture, selection and validation of CSA-binding phenotype**

