## Supporting information for "Novel monoclonal antibodies targeting distinct sites on placental binding *P. falciparum* antigen VAR2CSA synergistically enhance parasite phagocytosis"

**S1 Table: % afucosylation for mAbs**

| mAb | % Afucosylation | |
|---|---|---|
| | Complex Glycans | All Glycans |
| mPM1.13 | 1.1 | 26.5 |
| mPM1.14 | 0.7 | 24.5 |
| mPM1.15 | 0.9 | 30.8 |
| mPM1.16 | 1.2 | 27.7 |

### S2 Table: VAR2CSA mAb Sequences

| mAb name | IGHV Gene | IGHV homology (%) | IGJV Gene | IGDV Gene | CDR-H3 Length | CDR-H3 |
|---|---|---|---|---|---|---|
| mPM1.1 | Homsap IGHV3-64*07 F | 88.2 | Homsap IGHJ3*02 F | Homsap IGHD1-20*01 F | 16 | CTREGRYSWNNDDAFDIW |
| mPM1.2 | Homsap IGHV5-51*01 F | 92.4 | Homsap IGHJ3*01 F, or Homsap IGHJ3*02 F | Homsap IGHD2-21*01 F | 10 | CARLEDEAFDLW |
| mPM1.3 | Homsap IGHV3-64*01 F | 91.0 | Homsap IGHJ3*01 F | Homsap IGHD1-1*01 F | 16 | CTREGRYRWNDDDAFDVW |
| mPM1.4 | Homsap IGHV3-15*01 F | 90.8 | Homsap IGHJ5*02 F | Homsap IGHD3-3*01 F | 17 | CTKGGDDFWNDVYNWFDPW |
| mPM1.5 | Homsap IGHV1-69-2*01 F | 94.8 | Homsap IGHJ4*02 F | Homsap IGHD3-22*01 F | 15 | CATIPTYDSSGHYAGYW |
| mPM1.6 | Homsap IGHV3-7*01 F | 90.6 | Homsap IGHJ4*02 F | Homsap IGHD7-27*01 F | 10 | CVRGGWGDADHW |
| mPM1.7 | Homsap IGHV3-20*04 F | 82.3 | Homsap IGHJ4*02 F | Homsap IGHD3-22*01 F | 14 | CARDYYDSTGPLFDYW |
| mPM1.8 | Homsap IGHV3-20*05 F | 92.4 | Homsap IGHJ4*02 F | Homsap IGHD3-22*01 F | 14 | CARDYYDSTGPLYDYW |
| mPM1.9 | Homsap IGHV5-51*01 F | 94.1 | Homsap IGHJ3*02 F | Homsap IGHD5-18*02 ORF | 10 | CARLEDHAFDIW |
| mPM1.10 | Homsap IGHV3-64*07 F | 91.7 | Homsap IGHJ3*02 F | Homsap IGHD1-20*01 F | 16 | CTREGRYRWNNDDDFDIW |
| mPM1.11 | Homsap IGHV3-64*07 F | 88.5 | Homsap IGHJ3*02 F | Homsap IGHD1-1*01 F | 16 | CTREGRFSWNDDDAFDIW |
| mPM1.12 | Homsap IGHV4-39*02 F | 94.5 | Homsap IGHJ4*02 F | Homsap IGHD2-21*01 F | 10 | CTVVIFVSFDYW |
| mPM1.13 | Homsap IGHV5-51*01 F | 89.9 | Homsap IGHJ3*02 F | Homsap IGHD5-24*01 ORF | 10 | CARLKEHAFDIW |
| mPM1.14 | Homsap IGHV3-15*01 F | 93.2 | Homsap IGHJ5*02 F | Homsap IGHD3-3*01 F | 17 | CTKSGDDFWTDVYNWFDPW |
| mPM1.15 | Homsap IGHV4-39*01 F | 86.9 | Homsap IGHJ5*02 F | Homsap IGHD5-18*01 F | 10 | CATNTYREFDHW |
| mPM1.16 | Homsap IGHV3-64*07 F | 92.6 | Homsap IGHJ3*02 F | Homsap IGHD1-1*01 F | 16 | CTREGRFRWNDDDAFDIW |

| VDJ sequence |
|---|
| [illegible] |
| [illegible] |
| [illegible] |
| [illegible] |
| [illegible] |
| [illegible] |
| [illegible] |
| [illegible] |
| [illegible] |
| [illegible] |
| [illegible] |
| [illegible] |
| [illegible] |
| [illegible] |
| [illegible] |
| [illegible] |

**S2 Table: Summary of recombinant VAR2CSA proteins and mAb controls**

| Protein | Strain | Expression System | Provider or supplier | Reference |
|---|---|---|---|---|
| VAR2CSA | FCR3 | Mammalian (Expi293) | - | - |
| VAR2CSA | NF54 | Mammalian (Expi293) | - | - |
| VAR2CSA | M.Camp | Mammalian (Expi293) | Gift from Patrick Duffy and David Narum. NIH\| NIAID, USA | Renn et al, 2021 |
| VAR2CSA | HB3 | Mammalian (Expi293) | Gift from Patrick Duffy and David Narum. NIH\| NIAID, USA | Renn et al, 2021 |
| DBL1X-ID2a | FCR3 | *E. coli* | Gift from Morten Nielsen. University of Copenhagen, Denmark | |
| DBL3X | IT | | Gift from Patrick Duffy and David Narum. NIH\| NIAID, USA | |
| DBL4ε | M-711 | *E. coli* | Gift from Patrick Duffy and David Narum. NIH\| NIAID, USA | Doritchamou et al., 2016 |
| DBL5ε | 3D7 | *P. pastoris* | Gift from Patrick Duffy and David Narum. NIH\| NIAID, USA | Avril et al., 2011 |
| DBL6ε | IT4 | *P. pastoris* | Gift from Patrick Duffy and David Narum. NIH\| NIAID, USA | Avril et al., 2011 |
| IgG1 Isotype | - | Human cell Preparation | Sino Biological Cat#HG1K | - |
| PAM2.8 | - | Mammalian (Expi293) | Sequence was kindly passed on by Lars Hviid and Maria del Pilar Quintana Varon, University of Copenhagen, Denmark | Barfod et al., 2007 |

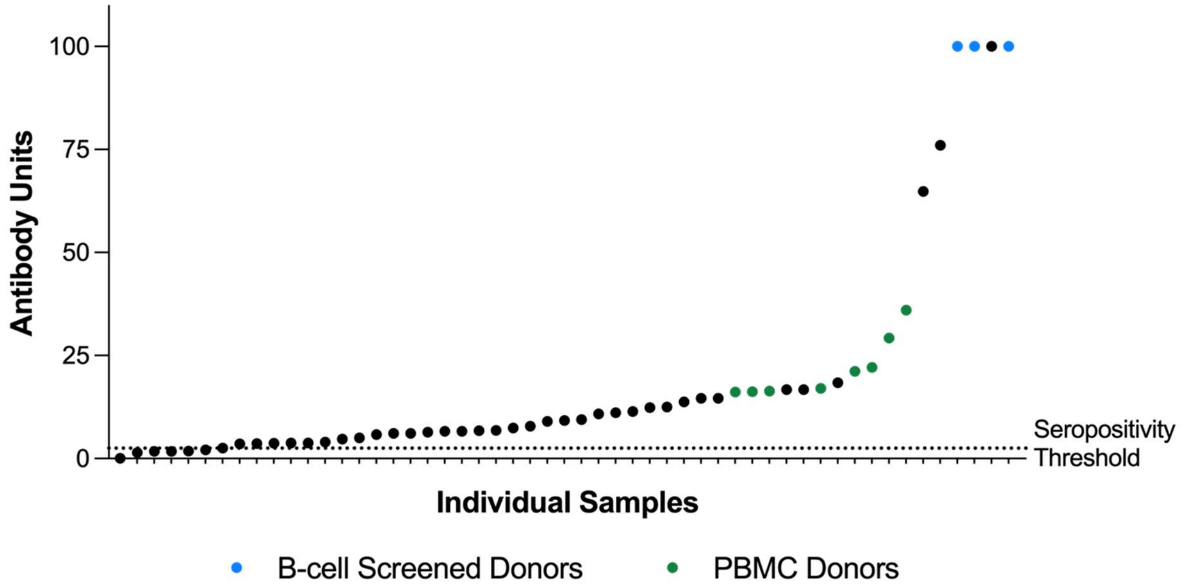


**S1 Figure: VAR2CSA antibody screen of participants of the SAPOT parent study.** Plasma samples from participants in the SAPOT study (n=53) were screened by ELISA to identify multigravidae with high VAR2CSA antibody levels. A subset of seropositive women (n=11) was recruited with informed consent, and their peripheral blood mononuclear cells (PBMCs) were collected for the present study. Three of the donors with the highest IgG levels to VAR2CSA were screened for B cells that cross-reacted with the NF54 and FCR3 VAR2CSA antigen baits to identify mAbs.

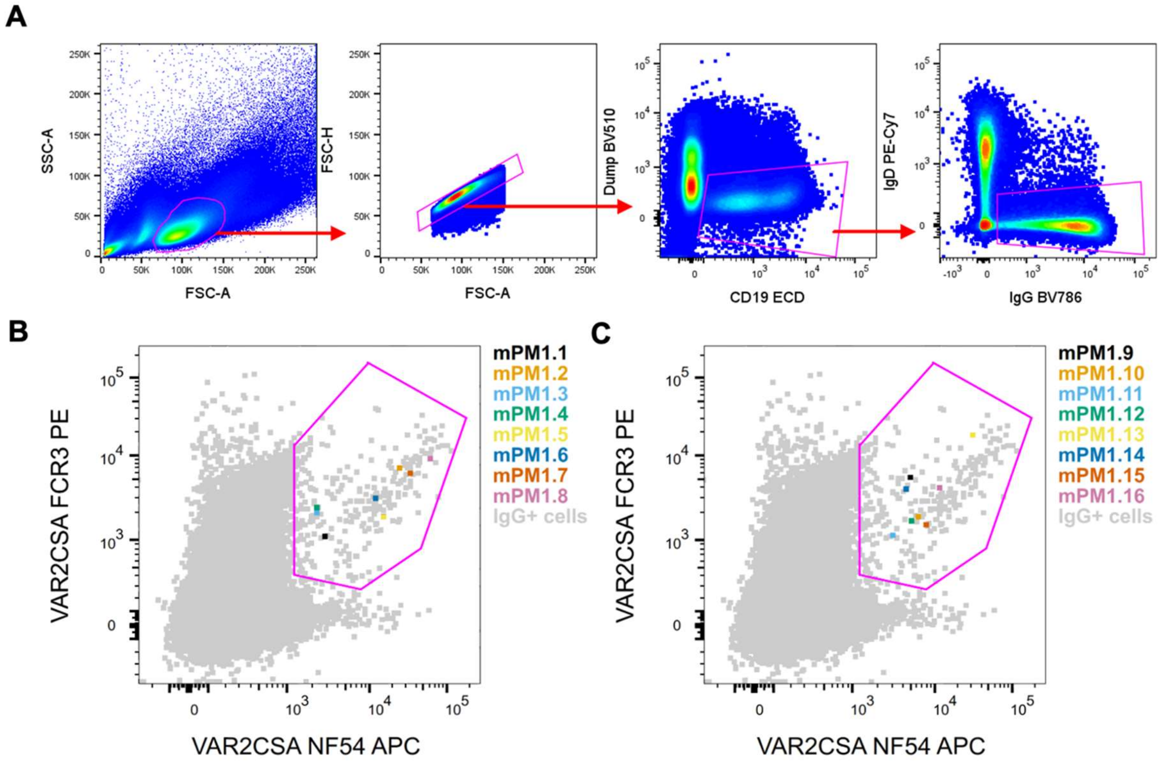


**S2 Figure: FACS gating and VAR2CSA probe staining with annotated B cells yielding successful mAbs.** (A) Gating used to identify IgG+ B cells from PBMCs. (B, C) VAR2CSA FCR3 and NF54 probe staining on IgG+ B cells. Sorted cells that yielded sequences for mPM1.1–mPM1.16 are indicated by colour according to the legends. The overall gate used to sort probe-positive cells is shown in pink.

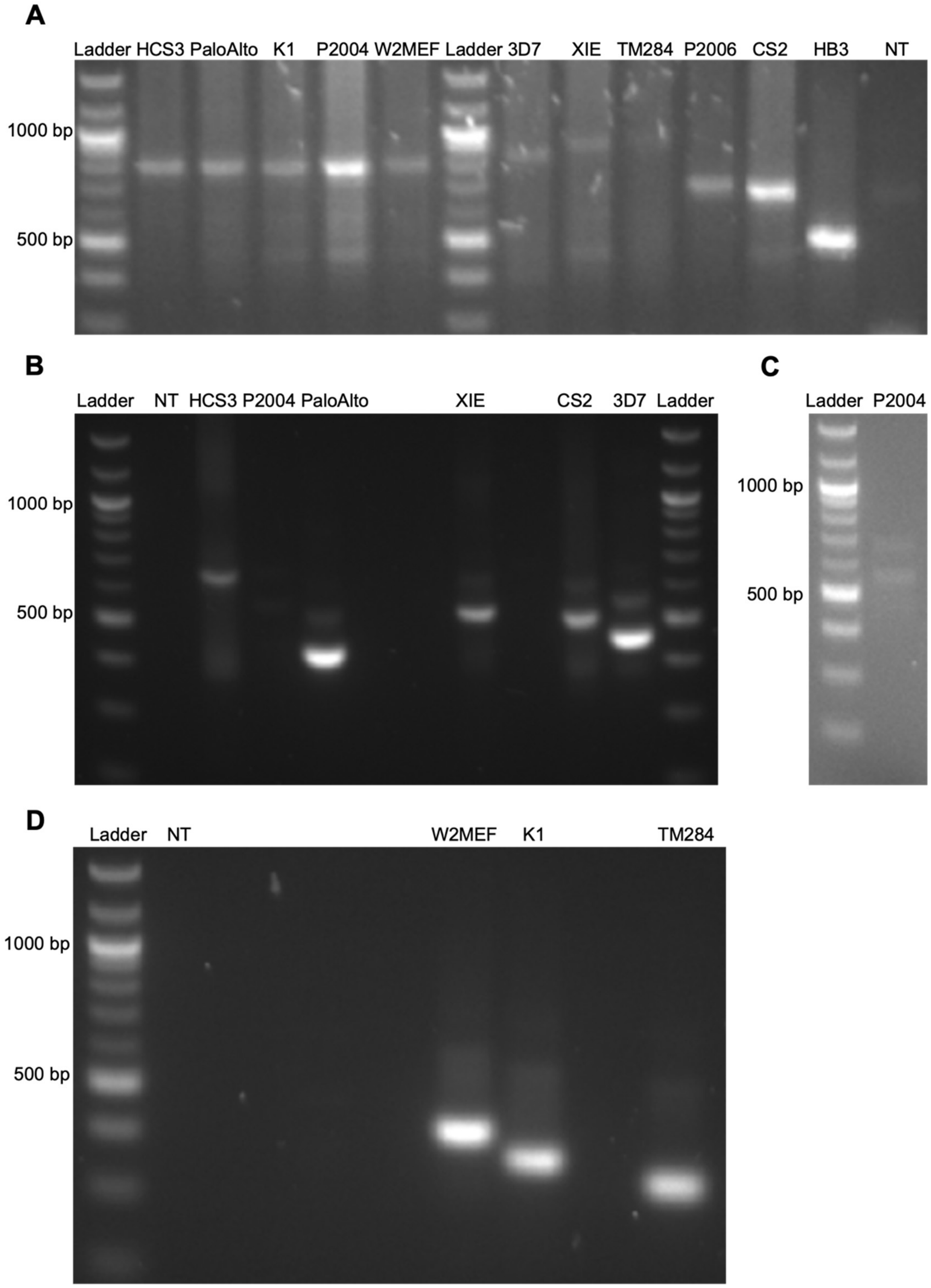


**S3 Figure: Genotyping of *glurp* and *msp2* of laboratory parasite isolates.** Genotyping of parasite isolates using primers specific to (A) *glurp, (B, C) msp2* of 3D7/IC allelic family and (D) *msp2* of FC27 allelic family demonstrated that all isolates were genotypically distinct. (C) A repeat gel for P2004 products from gel (B). Parasite isolates Palo Alto, W2MEF, and TM284 were not used for mAb characterisation, as Palo Alto could not be maintained in culture and W2MEF and TM284 did not express a CSA-binding phenotype. NT: no template, bp: base pair

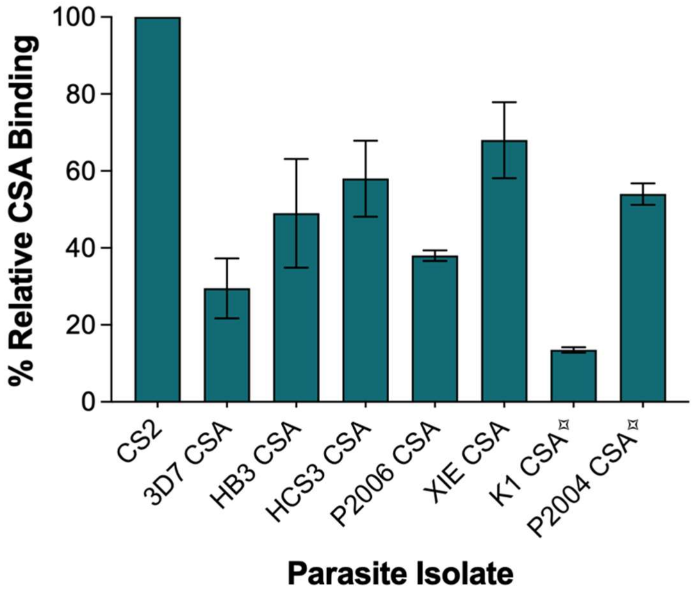


**S4 Figure: CSA-binding of heterologous parasite isolates.** Parasite isolates followed by -CSA were panned on immobilised CSA and/or BeWo cells to select for parasite phenotype. CSA binding of isolates to immobilised CSA was then measured as described by Dube *et al*. (2025), and binding levels were calculated relative to the CS2 isolate with constitutive expression of the CSA-binding phenotype. Bars represent mean ± SD from two individual experiments, except ✧, bars represent duplicates from an individual experiment.

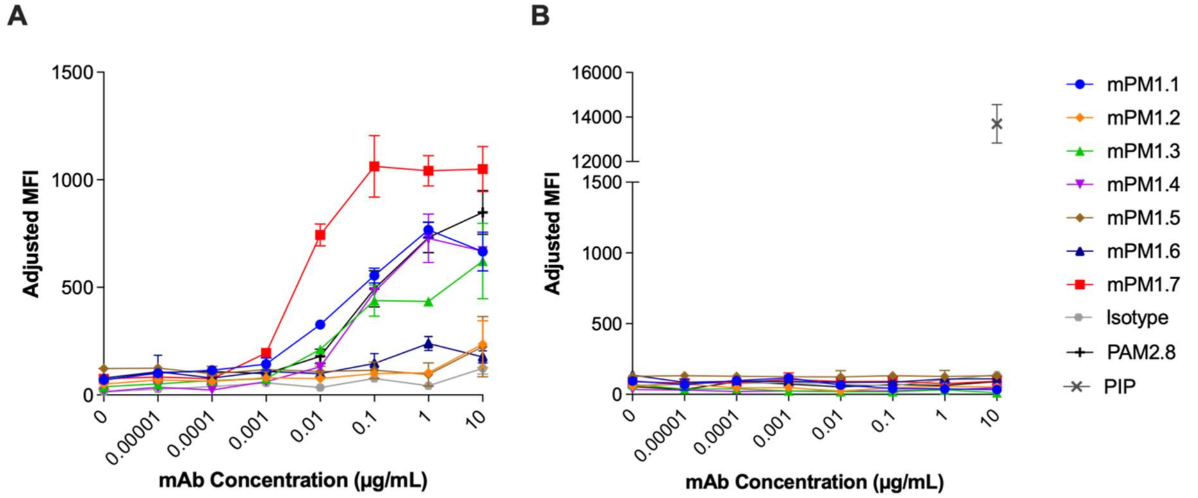


**S5 Figure: Representative mAb reactivity to infected erythrocytes (IEs) of CSA-binding and non-CSA-binding parasite isolates.** mAb reactivity to (A) CS2 parasite isolate with CSA binding phenotype and (B) E8B parasite isolate that does not predominantly bind CSA. Mean Fluorescence Intensity (MFI) is adjusted by subtracting the MFI of uninfected erythrocytes from that of IEs. Indicates mean and SD from experimental duplicates. PIP: Pooled Immune Plasma

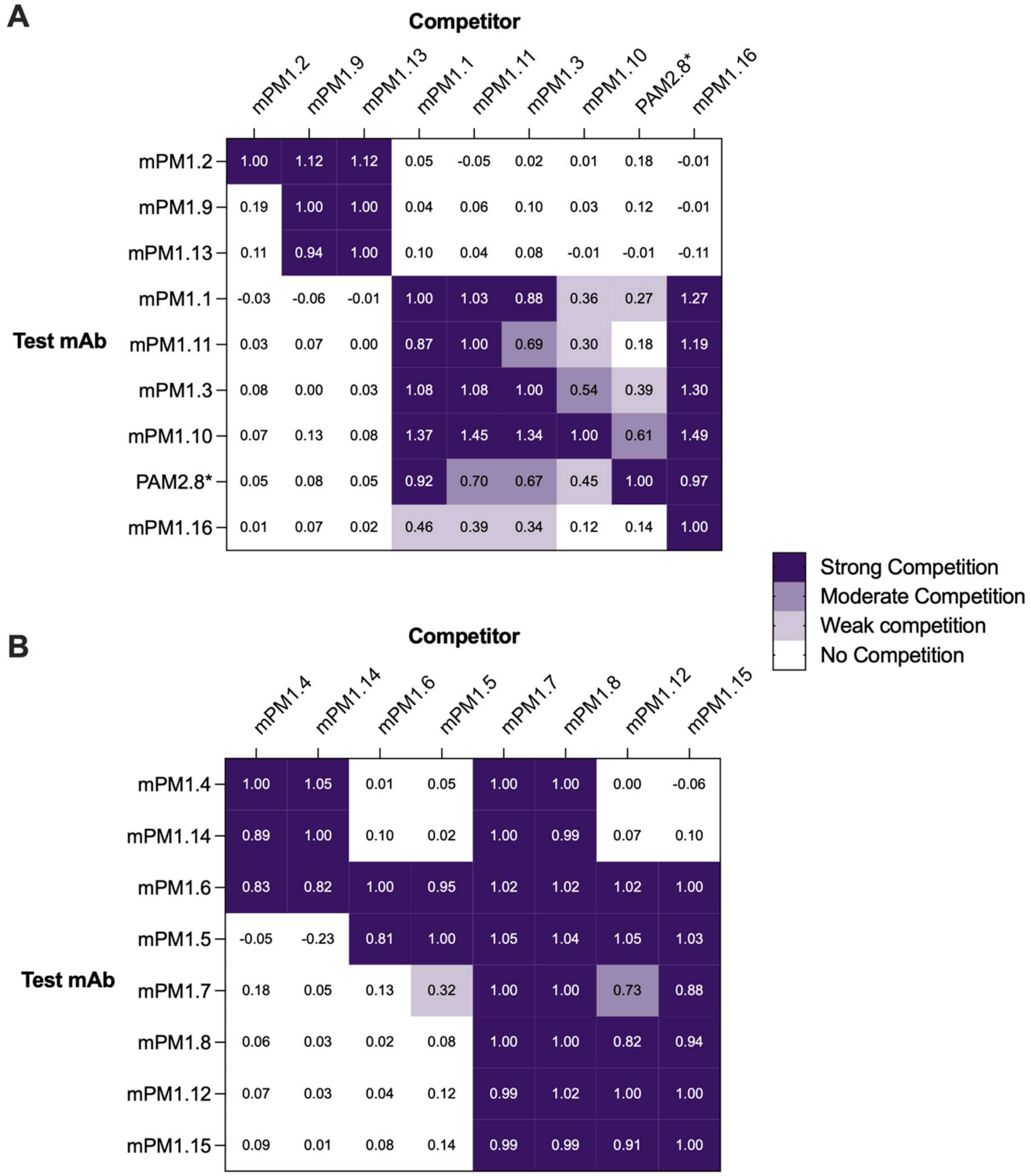


**S6 Figure: Competition matrices of mAbs targeting a single VAR2CSA domain.** The binding of a biotinylated test mAb was detected in the presence of an excess of competitor to assess whether two mAbs compete for a shared binding site. (A) Competition of mAbs targeting the DBL3X domain of VAR2CSA. (B) Competition of mAbs targeting the DBL5ε domain of VAR2CSA. Competition is calculated as optical density relative (rOD) to the self-competitor and is defined as strong (rOD ≥ 0.75), moderate (0.5 ≤ rOD < 0.75), weak (0.25 ≤ rOD < 0.5), or no competition (rOD ≤ 0.25). *PAM2.8 previously published mAb.

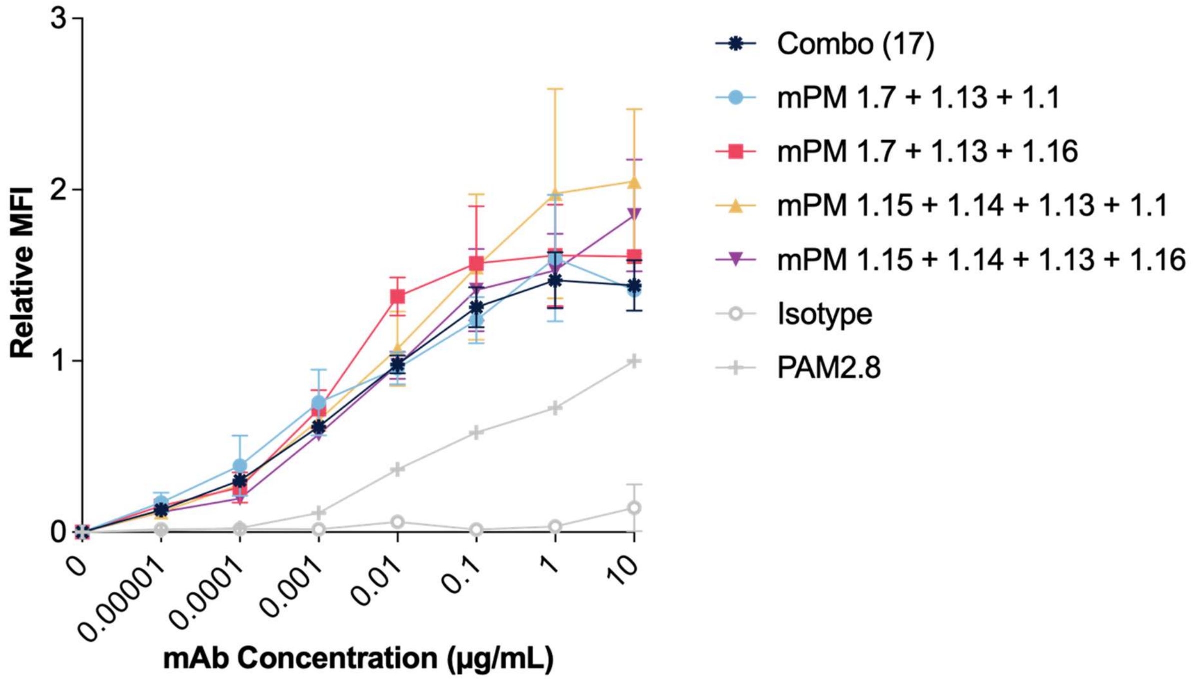


**S7 Figure: mAb cocktail reactivity to CS2 IEs.** The binding of mAb cocktail formulations to IEs was evaluated to assess whether opsonisation levels varied significantly across formulations. All conditions tested were at a total antibody concentration of 10 µg/mL. Mean fluorescence intensity (MFI) is calculated relative to PAM2.8 (previously published VAR2CSA mAb) at 10 µg/mL. The graph presents the mean ± SD from two experiments.

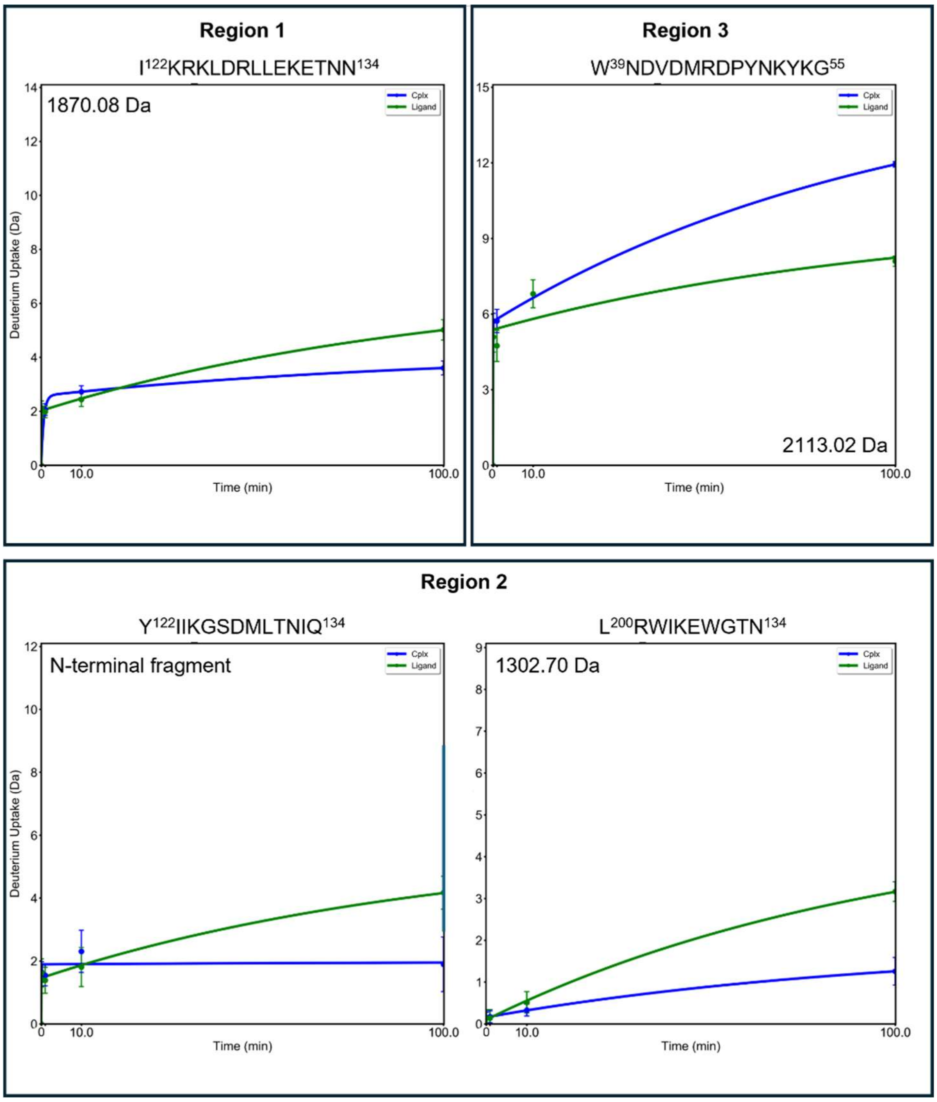


**S8 Figure: Uptake plots for selected peptides in the three noted regions of DBL5ε.** Peptides in regions 1 and 2 demonstrate significantly decreased deuterium uptake in the presence of mPM1.7, especially at long exposure, whilst region 3 demonstrates the inverse.

### S1 Methods: Mass spectrometric determination of afucosylation of mAbs

Purified mAbs were proteolysed overnight (17 hr) with Trypsin. Digested samples were analysed via Reverse Phase Liquid Chromatography-Mass Spectrometry (LC-MS/MS) under previously reported conditions (Ding. H et al., (Submitted); Falck & Wuhrer, 2024). In brief, 5 µL of peptide solution were loaded onto an Acclaim Pepmap nano-trap column (Dionex-C18, 100 Å, 75 µm × 2 cm), and then subsequently separated on a water-acetonitrile gradient through an Acclaim Pepmap RSLC analytical column (Dionex-C18, 100 Å, 75 µm × 50 cm). Mass spectrometry was performed on a Fusion Lumos Tribrid Mass Spectrometer (Thermo Fisher Scientific, Massachusetts, U.S.A) in a data-dependent manner, utilising dynamic exclusion of selected precursors for 60 seconds. MS data were analysed with

MSFragger in FragPipe (MSFragger version: 4.1) and the results were analysed in Skyline (Ver 23.1.0.455) (Kong et al., 2017; MacLean et al., 2010).

The % afucosylation was determined. 42 glycans were identified, of which 15 were definitively identified as complex glycans (G0F, G0, G0BF, G0B, G1F, G1, G1BF, G1B, G2FS, G2F, G2S, G2, G2BFS, G2BF, G2B). These complex glycans represented approximately 60%-70% of all observed glycans. The remaining glycans represented both high mannose glycans (6 glycans representing 23-25 %) and hybrid glycans (21 glycans). These results show less complex glycans than normally found on IgG, and may be a result of the expression system used.

**S2 Methods: Parasite culture, selection and validation of CSA-binding phenotype**

*P. falciparum* isolates were cultured and maintained as previously described in medium containing 5% human serum and 0.25% Albumax II (Yosaatmadja et al., 2008). The CSA-binding phenotype of laboratory isolates was enriched by panning on immobilised CSA. CSA (from bovine trachea; Sigma-Aldrich, 100 µg/mL in PBS) was immobilised on a petri dish in a humid box (16-18 hrs, 4 °C), washed (3x) with 1X PBS and blocked with 1% BSA (1hr, RT). Knobby trophozoites were selected through gelatin (0.75%) floatation, adjusted to 2% haematocrit in cytoadhesion medium (culture medium without $NaHCO_3$), added to the CSA-coated plate, and incubated in malaria gas mixture (1 hr, 37 °C) (Yosaatmadja et al., 2008). Unbound IEs were gently washed with RPMI-HEPES without $NaHCO_3$ until no unbound cells were observed under an inverted microscope. Bound cells were cultured in media with 3-5 drops of $O^+$ blood for 2-3 replication cycles until parasitaemia was observed under Giemsa-stained microscopy; cultures were then transferred to a new culture plate. This process was repeated to enrich cultures with CSA-binding parasites. Finally, a published colourimetric binding inhibition assay was adopted to measure CSA binding levels in enriched parasite isolates (Dube et al., 2025).

Parasite isolates Pf2004, Pf2006, and K1, which initially showed limited CSA binding, were panned on BeWo cells to enrich for CSA-binding parasites before selection was continued with immobilised CSA. BeWo cells were cultured in DMEM/F-12 medium containing L-glutamine and 2.40 g/L $NaHCO_3$ (Gibco), supplemented with 10% FBS and 1% penicillin/streptomycin. BeWo panning was performed as previously described (Yosaatmadja et al., 2008), with a few modifications: BeWo cells were seeded into a 6-well tissue culture plate for panning, unbound IEs were washed repeatedly until satisfactory under an inverted microscope, and cultured bound IEs overnight in parasite culture medium with 2-3 drops of

$O^+$ blood, before transferring ring-stage IEs into a new 6-well plate for continued culture. The selection process was repeated three times to obtain a higher number of CSA-binding parasites.
